# Subthreshold repertoire and threshold dynamics of midbrain dopamine neuron firing *in vivo*

**DOI:** 10.1101/2020.04.06.028829

**Authors:** Kanako Otomo, Jessica Perkins, Anand Kulkarni, Strahinja Stojanovic, Jochen Roeper, Carlos A. Paladini

## Abstract

The firing pattern of ventral midbrain dopamine neurons is controlled by afferent and intrinsic activity to generate prediction error signals that are essential for reward-based learning. Given the absence of intracellular *in vivo* recordings in the last three decades, the subthreshold membrane potential events that cause changes in dopamine neuron firing patterns remain unknown. By establishing stable *in vivo* whole-cell recordings of >100 spontaneously active midbrain dopamine neurons in anaesthetized mice, we identified the repertoire of subthreshold membrane potential signatures associated with distinct *in vivo* firing patterns. We demonstrate that dopamine neuron *in vivo* activity deviates from a single spike pacemaker pattern by eliciting transient increases in firing rate generated by at least two diametrically opposing biophysical mechanisms: a transient depolarization resulting in high frequency plateau bursts associated with a reactive, depolarizing shift in action potential threshold; and a prolonged hyperpolarization preceding slower rebound bursts characterized by a predictive, hyperpolarizing shift in action potential threshold. Our findings therefore illustrate a framework for the biophysical implementation of prediction error and sensory cue coding in dopamine neurons by tuning action potential threshold dynamics.

---

The midbrain dopamine system is necessary for essential brain functions related to reward-based learning, motivation, action, and cognition (Athalye et al., 2020; Gershman and Uchida, 2019; Langdon et al., 2018; Schultz, 2015). Loss or dysregulation of dopamine neuron subpopulations located in the substantia nigra pars compacta (SNc) and ventral tegmental area (VTA) perturbs crucial circuits leading to major brain disorders, including Parkinson’s disease (Obeso et al., 2017), schizophrenia (McCutcheon et al., 2019), depression (Grace, 2016), and addiction (Volkow et al., 2017). However, describing the dynamics of subthreshold ionic conductances that cause dopamine neuron spiking activity *in vivo* has remained a challenge for neurophysiologists.

Single-unit extracellular recording studies from non-human primates and rodents have provided major insights into the computational roles of dopamine neuron firing, most prominently in the context of reward-based learning (Hikosaka et al., 2018; Schultz, 2015, 2016). Dopamine neurons signal prediction errors by transient, sub-second changes in their firing rates. A recent study implied that prediction error coding in dopamine neurons might occur in a distributed manner, thereby creating a range from “optimistic” to “pessimistic” cellular coding schemes (Dabney et al., 2020). For many years, transient cue- or reward-associated increases in firing rate – from a low tonic background frequency (about 1-8 Hz) into the beta and gamma range (15-30 Hz and 40-100 Hz, respectively) – were thought to encode positive reward prediction errors (RPEs), while transient reductions or pauses in baseline firing were believed to encode negative RPEs, induced for instance by the omission of an expected reward. However, increasing experimental evidence points to a multitude of other dopamine functions, such as salience and novelty, action control (Coddington and Dudman, 2018; Jin and Costa, 2010) as well as aversion (Menegas et al., 2018), which are in line with the emerging molecular (Heymann et al., 2020; Poulin et al., 2020), cellular, and anatomical diversity of the midbrain dopamine system (Beier et al., 2019; Beier et al., 2015; Farassat et al., 2019; Lammel et al., 2008; Lammel et al., 2012; Morales and Margolis, 2017; Watabe-Uchida et al., 2012). Even within canonical RPE signaling, the interpretation of the transient burst has become more complex. It is currently divided into a short-latency component, which signals sensory characteristics of the stimulus (e.g. intensity), and a long-latency component, which represents predictive properties of the stimulus (Schultz, 2016). In addition, recent progress has been made in characterizing the underlying arithmetic of single dopamine neurons in the context of RPE signaling (Eshel et al., 2015). It is reasonable to assume that dopamine neurons *in vivo* integrate various heterogeneous afferents throughout the brain (Tian et al., 2016) to tune intrinsic pacemaker activity and generate burst-pause firing patterns, which in turn lead to temporally-resolved, extracellular dopamine concentration transients in target regions (Beeler and Kisbye Dreyer, 2019). In addition, mechanisms of dopamine firing synchronization, as well as presynaptic control, are likely to shape axonal dopamine release (Condon et al., 2019). Therefore, dopamine neurons are not simply leaky integrators of excitatory and inhibitory synaptic inputs, but display a rich and diverse repertoire of intrinsic excitability, including biophysical mechanisms for autonomic pacemaker firing.

However, how inputs are integrated *in vivo* at the cellular level has mostly remained unknown for many decades due to the lack of available tools. While the *in vivo* extracellular recording method does record spontaneous firing patterns generated by dopamine neurons, it is blind to the causal subthreshold processes that lead to the apparent timed spiking patterns. Intracellular recordings are unique in their ability to directly monitor and manipulate membrane potentials with full temporal resolution. Yet, the only available intracellular *in vivo* data on dopamine neurons came from the pioneering studies by Grace & Bunney in the early eighties (Grace and Bunney, 1980, 1983a, b, 1984a, b). Here, we provide the first large dataset in nearly four decades on the subthreshold membrane potential behavior *in vivo* associated with firing patterns such as pacemaking and bursting. We established a robust method for deep *in vivo* patch-clamp recordings of dopamine neurons in anaesthetized, adult C57BL/6 mice. Our dataset comprises 112 spontaneously active, neurobiotin-labeled and immunocytochemically-identified dopamine neurons, which were mapped to their respective anatomical positions within the SNc and VTA. Unexpected by decades of *in vivo* extracellular recordings, we identified two distinct biophysical subthreshold signatures that drive transient burst firing. We demonstrate that pacemaking dopamine neurons *in vivo* have the ability to tightly control their excitability, as monitored by their action potential thresholds and interspike membrane potentials, and that opposing mechanisms underlying burst discharge are characterized by predictive and reactive shifts in action potential thresholds, and baseline membrane potentials. Thus, our study provides a framework for mechanistic studies into prediction error coding of dopamine neurons in awake and behaving animals.

## Results

Using our newly established deep *in vivo* patch-clamp recording technique (Fig. 1A and 1B), we have obtained *in vivo* whole-cell recordings from 112 spontaneously firing, identified midbrain dopamine neurons in isoflurane-anaesthetized, adult C57BL/6 mice (n = 112, N = 82; Fig. 1). All neurons included in this study were filled with neurobiotin or biocytin and had been successfully recovered for anatomical mapping and post-hoc identification as dopaminergic using tyrosine hydroxylase (TH) immunohistochemistry (Fig. 1C). While we densely sampled dopamine neurons in the rostral parabrachial VTA and rostro-medial SNc, we had less coverage in lateral aspects of the SNc and in more caudal ventro-medial, paranigral VTA regions (Fig. 1D). Due to the unbiased nature of *in vivo* patch-clamping, we also encountered TH-negative, electrically silent cells with membrane potentials below −50 mV as well as TH-negative, fast firing cells (>10 Hz) with narrow action potentials (data not shown). Moreover, in addition to the 112 spontaneously active dopamine neurons included in this study, we also observed 2 electrically silent TH-positive cells that would fire in response to depolarizing current injection, and 2 other TH-positive neurons that fired less than one spike per minute (data not shown). Under isoflurane anesthesia, the technical and biological stability of recordings in the whole-cell configuration were not a major limiting factor (Supplemental Figure 1). In addition, membrane potential responses to hyperpolarizing current injections were preserved *in vivo* and displayed a diversity reminiscent of previous *in vitro* studies (Supplemental Figure 2) (Lammel et al., 2008). Figure 1B shows a typical *in vivo* patch-clamp recording of an identified midbrain dopamine neuron. The recording presented on an expanded timescale displayed episodes of overshooting spikes interspersed by pauses where the membrane potential hyperpolarizes below the threshold for action potentials (Fig. 1B, bottom trace). Post-hoc immunohistochemistry identified this cell to be a dopamine neuron localized in the dorso-rostral VTA (Fig. 1C and 1D, red circle). Our dataset includes dopamine neurons recorded in male and female mice from two different C57BL/6 substrains (C57BL/6N and C57BL/6J), using two different versions of the pipette solution with a difference in estimated free calcium concentrations of 40 nM and 80 nM. As there were no significant differences in the mean firing rate (FR) and variabilities (quantified as coefficient of variation, CV) between these data (see Supplemental Table 1), we pooled the data. Figures 1E and 1F show the mean firing rates and CVs of spontaneous firing activity from the 112 dopamine neurons that passed our quality criteria. The input resistance was calculated for a subset of these neurons, which had a median of about 300 MΩ (Median (InterQuartile Range) = 295.16 (228.14 to 349.8) MΩ; n = 42, N = 36; Fig. 1G), an order of magnitude higher than previously reported values using sharp microelectrodes (Grace and Bunney, 1980 and 1983a). This high *in vivo* input resistance indicates that differences in intrinsic excitability of dopamine neurons are less likely to be shunted in comparison to other types of central neurons (Berg et al., 2007; El Boustani et al., 2007). The instantaneous firing rate range for each identified dopamine neuron plotted in Figure 1H illustrates a wide range of firing frequencies within our recordings.

**Figure 1.**
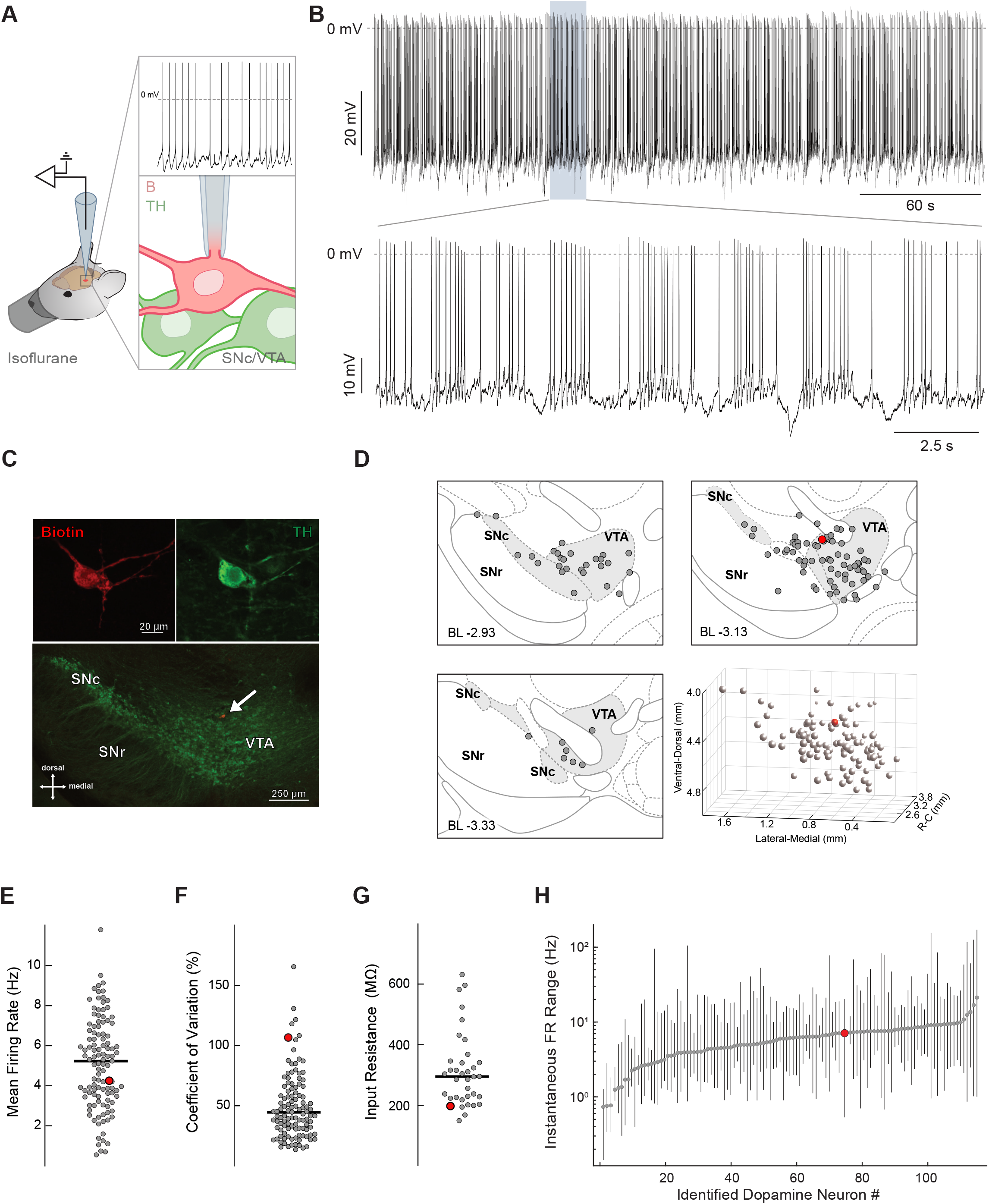
*In vivo* whole cell recordings of identified ventral mesencephalon dopamine neurons. **A.** Schematic representation of the experimental setup. **B.** Representative recording of spontaneous *in vivo* electrical activity from a whole-cell-recorded and neurochemically-identified dopamine neuron in the ventral tegmental area (VTA) of an adult C57Bl/6J mouse under isoflurane anesthesia. The upper trace displays action potentials with overshoot during a stable recording for > 5 minutes. The higher temporal resolution of the electrical activity in the lower trace allows visual identification of subthreshold events such as large hyperpolarizations and synaptic events. **C.** Immunocytochemical identification demonstrates the recorded neuron (top panel, filled with Biotin; red) is positive for tyrosine hydroxylase (TH; green). Lower magnification of the immunocytochemical image (lower panel) locates the recorded neuron (arrow) within the dorsal region of the VTA. **D**. Depiction shows the location of the recorded neuron (large red dot) in relation to all the recorded neurons in this study (gray dots) displayed at three coronal planes. Bottom right panel is a 3D representation of the example cell location (red sphere) in relation to all recorded cells (gray spheres). **E, F.** Plot of the mean firing rate (E) and coefficient of variation (CV) for interspike intervals (F) for each neuron presented in this study (horizontal line = median). **G.** Plot of the input resistance for each neuron where resistance was measured (horizontal line = median). **H.** Log-scale plot of the instantaneous firing rate, and its full range, for each neuron presented in this study sorted in the order of increasing median rate. **D-H.** The large red symbol in all plots represents the example neuron illustrated in B and C.

We subsequently focused on a systematic investigation of the subthreshold activity associated with tonic and phasic firing patterns in the intact brain. We found that a majority of neurons fired in a single spiking pattern, similar to our previous *in vivo* extracellular studies of identified dopamine neurons (Fig. 2) (Farassat et al., 2019; Schiemann et al., 2012). A representative recording, and the immunohistochemistry of an *in vivo* pacemaking dopamine neuron located in the VTA, is shown in Figures 2A and 2B. The trace with higher temporal resolution (Fig. 2A, bottom trace) demonstrates the stability of the *in vivo* pacemaker firing mode. To capture the subthreshold characteristics of *in vivo* pacemaking, we plotted the distribution of subthreshold membrane potentials for interspike interval (ISI) membrane potential minima (V_min_), i.e. the most hyperpolarized membrane potential between two action potentials (Fig. 2A, blue circles, bottom trace; Fig. 2C), as well as the distribution of subthreshold membrane maxima (V_thr_), i.e. the action potential thresholds (Fig. 2A, green circles, bottom trace; Fig. 2D). Both V_min_ and V_thr_ exhibited a narrow unimodal distribution, quantified by their respective unimodality indices (bimodality coefficient (BC): V_min_ = 0.35 ± 0.03; V_thr_ = 0.34 ± 0.05; n = 22, N = 21) and the standard deviations (SD) of the Gaussian fits (V_min_ mean = −42.8 ± 5.48 mV, SD = 1.39 ± 0.59 mV; V_thr_ mean = −30.59 ± 3.53 mV, SD = 0.88 ± 0.2 mV; Figures 2C and 2D). This demonstrates the capacity of *in vivo* dopamine neurons to maintain their stability in firing, similar to what is observed *in vitro,* even when embedded within active circuits. To examine a potential pattern between subthreshold and firing properties, V_min_ and V_thr_ are plotted against their respective ISI durations (Fig. 2E). While there is more variability in the subthreshold membrane potential minima as compared to the maxima, the respective ISIs of both plots show a high degree of stable spiking in pacemaking dopamine neurons. Next, we utilized event-triggered averaging to obtain a visual representation of a stereotypical subthreshold membrane potential signature of *in vivo* pacemaking activity (Fig. 2F). The averaged trace (Fig. 2F, bold trace) portrays a low-variance subthreshold signature of a ~200 ms gradual, ramp-like membrane potential depolarization of about 10 mV until reaching the firing threshold. Figure 2G shows the firing frequency distribution of this example cell. The frequency distribution of the entire population of dopamine neurons identified by their subthreshold properties to discharge in *in vivo* pacemaker mode is shown in Figure 2H.

**Figure 2.**
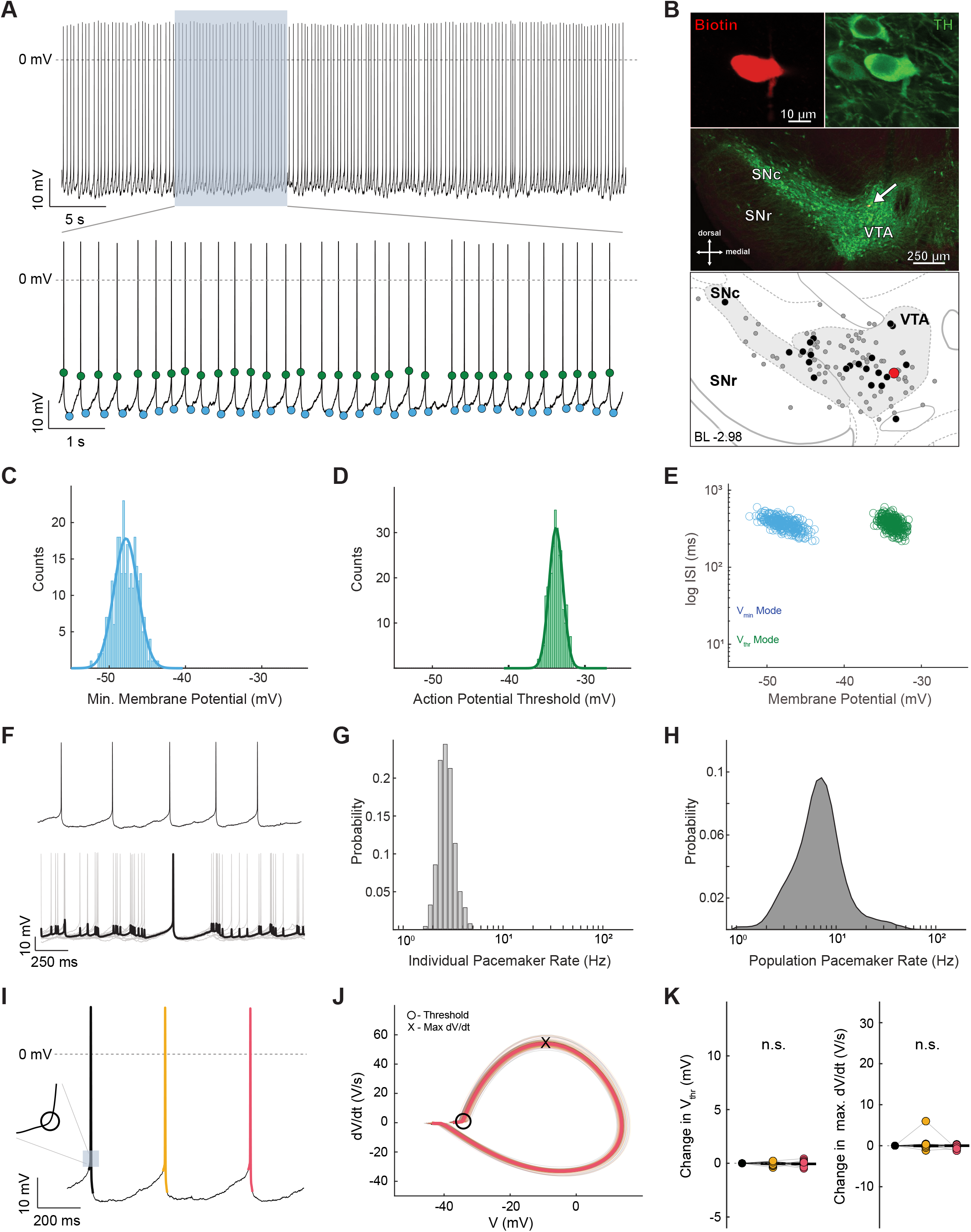
Pacemaker pattern is characterized by a stable and narrow subthreshold membrane potential dynamic range. **A**. Representative recording of spontaneous *in vivo* electrical activity from a whole-cell-recorded and neurochemically-identified midbrain dopamine neuron in the VTA. The upper trace displays action potentials with overshoot during a stable recording for > 60 s. The higher temporal resolution of the electrical activity in the lower trace allows visual identification of subthreshold events. Blue dots represent the most hyperpolarized membrane potential (Minima) reached in between every action potential. Green dots represent the most depolarized membrane potential (Maxima; action potential threshold) reached in between every action potential. **B.** Immunocytochemical identification demonstrates the recorded neuron (red) as TH-positive (green). Lower magnification of the image locates the recorded neuron (arrow) within the dorsal region of the VTA. The depiction shows the location of the recorded neuron (large red dot) in relation to all the recorded neurons in this study (gray and black dots). Black dots represent the locations of all the neurons that fired in a pacemaker pattern. **C.** Distribution of all subthreshold voltage minima (V_min_) from the cell in A-B, demonstrating a unimodal distribution (blue line = single Gaussian fit). **D.** Distribution of all subthreshold voltage maxima (V_thr_) from the cell in A-B, demonstrating a unimodal distribution (green line = single Gaussian fit). **E.** Scatter plot of the interspike interval (ISI) versus V_min_ (blue) and V_thr_ (green). Longer duration ISIs are correlated with more hyperpolarized voltage minima. **F.** High temporal resolution of pacemaker firing pattern (upper panel), and a mean spike-triggered pacemaker pattern (black trace in lower panel) with overlaid (gray) traces. **G.** Histogram of the instantaneous firing rate for the cell in A-B. **H.** Histogram of instantaneous firing rate for all cells firing in a pacemaker pattern. **I.** Example trace of a cell firing in a pacemaker pattern with three action potentials in order from black, then orange, then red. Inset illustrates where the threshold occurs for the black action potential. **J.** The phase plots for each action potential are displayed in the same color scheme as in I (black, orange, red; plots are nearly completely overlapping). **K.** Summary data of the changes in threshold voltage (left) and maximal depolarization rate (Max dV/dt; right) across the sequence of three action potentials for all cells firing in a pacemaker pattern.

To gain more mechanistic insight, we analyzed the phase plots of action potentials in pacemaking firing sequences. Figure 2I illustrates an example of a three-action potential sequence (black - orange - red) during pacemaker firing. A phase plot for all spikes occurring during pacemaker firing from one dopamine neuron is shown in Figure 2J. Note the large degree of overlap throughout the different phases of the action potentials, from threshold to upstroke and overshoot, as well as during repolarization and afterhyperpolarization. Quantitative analysis of all dopamine neurons discharging in this *in vivo* pacemaker mode (n = 22, N = 21) confirmed a high degree of stability and low variability that extended from the subthreshold domain to the action potential. Indeed, we did not observe significant changes in threshold voltage, maximal depolarization (Max dV/dt), or repolarization (Min dV/dt) speeds across the sequence of three action potentials (Δ V_thr_: spike 1 vs spike 2 = −0.03 (−0.09 to 0.17) mV, spike 2 vs 3 = −0.05 (−0.1 to 0.08) mV; Δ max dV/dt: spike 1 vs 2 = −0.04 (−0.26 to 0.31) V/s, spike 2 vs 3 = −0.03 (−0.3 to 0.17) V/s; p = 0.59, 0.79, 0.59, and 0.52 for all comparisons; n = 22, N = 21; Fig. 2K and Supplemental Table 2).

We subsequently studied transient high frequency firing episodes in SNc/VTA dopamine neurons, usually termed “bursts”. In contrast to cue- and reward-triggered bursts in awake animals, *in vivo* burst firing in anaesthetized animals has no clear behavioral correlate, and thus seem to occur spontaneously. Moreover, bursts do not only manifest as single events, but often occur in patterns, with some dopamine neurons being locked in a repetitive burst firing mode (Paladini and Tepper, 1999; Schiemann et al., 2012). We therefore focused on the dopamine neurons locked in burst modes, which enabled for the first time the unbiased identification and classification of burst-related subthreshold membrane potential signatures *in vivo* without relying on ISI-based heuristics (Bingmer et al., 2011; Grace and Bunney, 1984a; Ko et al., 2012). To our surprise, spontaneous *in vivo* bursting of identified dopamine neurons occurred in association with two diametrically opposing subthreshold membrane potential signatures (compare Figures 3 and 4). One of the two burst modes, illustrated in Figure 3, is characterized by a transient, high frequency discharge preceded by a prolonged and pronounced membrane potential hyperpolarization (Fig. 3). Accordingly, we termed this mode *rebound burst* firing. In contrast to the example pacemaking dopamine neuron we showed in Figure 2, the distribution of V_min_ was best described as bimodal Gaussian distributions (Mode 1 and Mode 2; Fig. 3C). This was quantified by calculating the bimodality coefficient of V_min_ distributions for all dopamine neurons with rebound bursting (V_min_ BC = 0.64 ± 0.14; Mode 1 V_min_ mean = −46.37 ± 3.88 mV, SD = 2.64 ± 1.24 mV; Mode 2 V_min_ mean = −39.39 ± 4.34 mV, SD = 1.49 ± 0.45; n = 20, N = 19). Importantly, when plotting ISI durations against V_min_, the more strongly-hyperpolarized membrane potentials (Fig. 3C, Mode 1) were almost exclusively associated with longer ISI durations in the range of 640-1120 ms, forming an independent cluster in the plot (Fig. 3E, dark blue circles), whereas the more depolarized V_min_ were associated with a wide range of shorter ISI durations (Fig. 3E, light blue circles). In contrast, the distribution of V_thr_ in rebound burst dopamine neurons was unimodal (Fig. 3D and 3E, green circles) and did not show a similar segregation with corresponding ISI durations (V_thr_ mean = −30.03 ± 3.01 mV, SD = 1.42 ± 0.49 mV). We applied event-triggered averaging to spikes that immediately followed the Mode 1 V_min_ events to reveal the low-variance subthreshold membrane potential signature of rebound bursting, (Fig. 3F, bottom trace). The rebound bursting subthreshold signature was characterized by a long (559.03 ± 218.96 ms, n = 20, N = 19) and curved trajectory of initial membrane hyperpolarization (8.65 ± 3.03 mV hyperpolarization to a minimum of −48.04 ± 3.95 mV, n = 20, N = 19) followed by membrane depolarization back to threshold. This rebound burst mode was associated with bursting in the beta frequency range, but also allowed lower frequencies. In contrast, event-triggered averaging of spikes following the Mode 2 V_min_ events did not result in a stereotypical low-variability subthreshold waveform (data not shown). Thus, the frequency distribution associated with the rebound burst significantly widened the dynamic range of dopamine neurons compared to pacemaking (Median pacemaker firing rate: 6.3 (4.43 to 8.27) Hz, p = 2.43 x 10^−262^ versus rebound burst firing rate), both on the level of individual cells and, in particular, on the level of dopamine neuron populations (compare Fig. 2H and Fig. 3H). This is in line with the recent idea of a distributed population code for dopamine neurons (Dabney et al., 2020).

**Figure 3.**
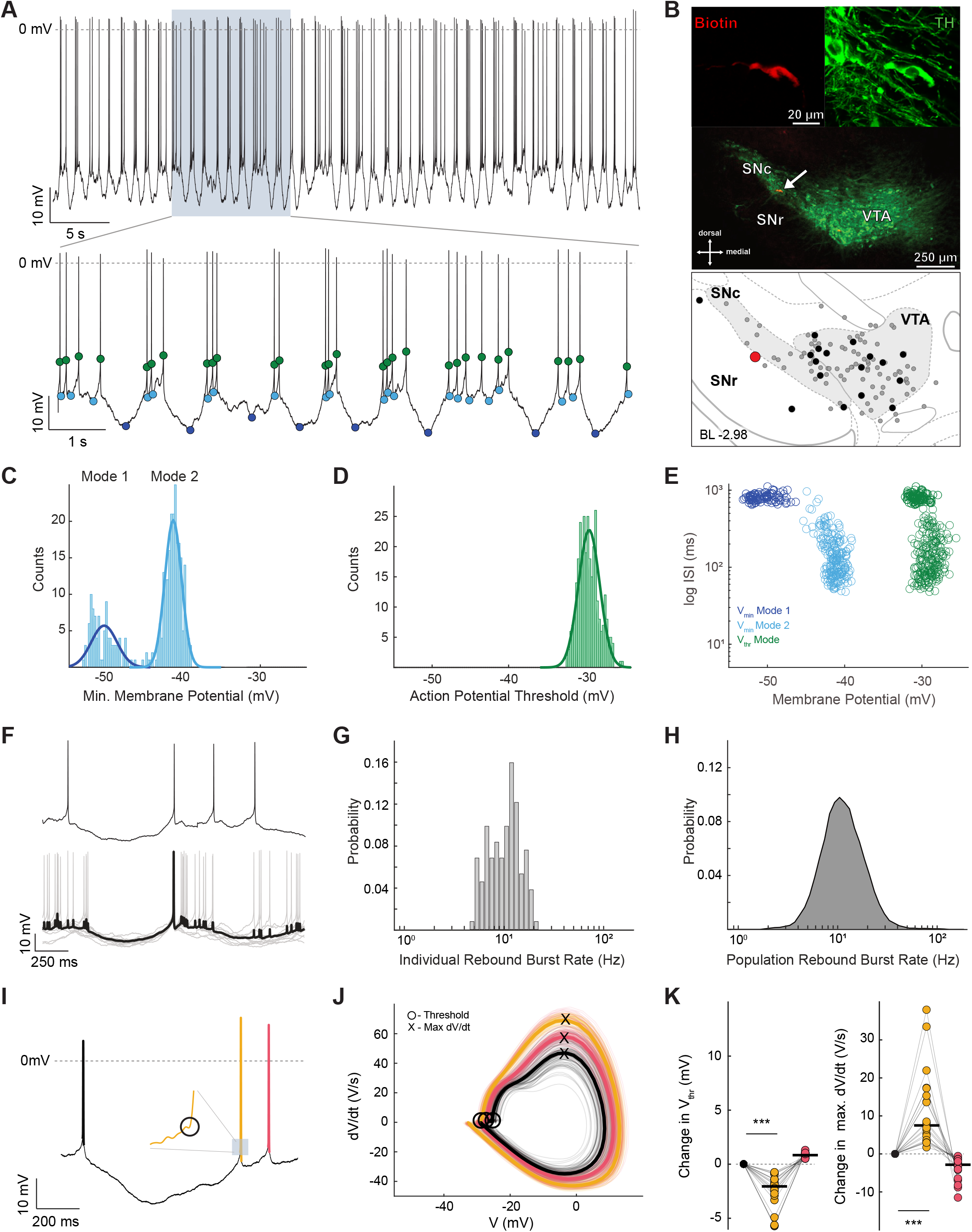
Rebound burst is characterized by a large hyperpolarization and a hyperpolarizing action potential threshold. **A**. Representative recording of an identified dopamine neuron in the SNc. The upper trace displays action potentials with overshoot during a stable recording for > 60 s. The higher temporal resolution in the lower trace allows visual identification of subthreshold events. Blue dots represent the Minima reached in between every action potential. The dark blue dots represent the minima of large hyperpolarizations. Green dots represent the Maxima (i.e. action potential threshold). **B.** Immunocytochemical identification (top panel) demonstrates the recorded neuron (red) as TH-positive (green). Lower magnification image (middle panel) locates the neuron (arrow) within the SNc. The depiction (bottom panel) shows the location of the recorded neuron (large red dot) in relation to all the recorded neurons (gray and black dots). Black dots represent the locations of all neurons that fired in a rebound burst pattern. **C.** Distribution of subthreshold voltage minima (V_min_) from the cell in A-B, demonstrating a bimodal distribution (dark blue line = Mode 1 Gaussian fit; light blue line = Mode 2 Gaussian fit). **D.** Distribution of subthreshold voltage maxima (V_thr_) from the cell in A-B, demonstrating a unimodal distribution (green line = single Gaussian fit). **E.** Scatter Plot of ISI versus V_min_ and V_thr_. The dark blue cluster is associated with more hyperpolarized voltage membrane potentials and longer duration ISIs. **F.** High temporal resolution of a rebound burst (upper panel), and event-triggered rebound burst average (lower panel, black trace) and overlaid (gray) traces. **G.** Histogram of the cell firing rate associated with rebound bursts. **H.** Histogram of firing associated with rebound bursts for all cells that fired in a rebound burst pattern. **I.** Example trace of a cell firing a rebound burst with three action potentials in order from black prior to the burst, then orange beginning the burst, and red. Inset illustrates where the hyperpolarized threshold occurs for the action potential beginning the burst (orange). **J.** The phase plots for each action potential are displayed in the same color scheme as in I. **K.** Summary data of the changes in threshold voltage (left) with significant hyperpolarization in threshold of the first action potential of the burst (orange), and change in maximal depolarization rate (Max dV/dt; middle) with significant increase in Max dV/dt, across the sequence of three action potentials for all cells firing in a rebound burst pattern.

The phase plot of the rebound burst action potentials shows different properties compared to that of pacemaking (Fig. 3I-3K). In particular, the first action potential of the rebound burst series (Fig. 3I, orange) had a significantly more hyperpolarized threshold associated concurrently with an accelerated speed of depolarization (Max dV/dt; Fig. 3J and 3K). These changes are consistent with an enhanced excitability, e.g. by increasing the number of available voltage-gated sodium channels set by the membrane hyperpolarization that precedes the onset of the rebound burst. Once within the rebound burst, the threshold and depolarization speed return close to pre-burst baseline already with the next spike (Fig. 3J; Δ V_thr_: spike 1 vs spike 2 = −2.06 (−3.12 to −1.4) mV, spike 2 vs 3 = 0.82 (0.63 to 1.03) mV; Δ max dV/dt: spike 1 vs 2 = 7.49 (3.49 to 16.3) V/s, spike 2 vs 3 = −2.9 (−5.32 to −1.74) V/s; p = 8.01 x 10^−9^, 6.8 x 10^−8^, p = 8.01 x 10^−9^, 6.8 x 10^−8^ for all comparisons; n = 20, N = 19; Fig. 3K and Supplemental Table 2). Thus, the observed threshold dynamics of rebound bursts in dopamine neurons occur at burst onset.

In comparison to the rebound burst mode, the second burst mode has qualitatively different subthreshold properties that were exclusively associated with higher frequency bursting (Fig. 4). This mode of bursting was characterized by a very short transient high frequency event in the gamma range and was identified by with a large positive shift in the V_thr_. As these spikes occurred on a depolarized “plateau” of the membrane potential, we termed this discharge pattern *plateau bursts*. An example trace and its corresponding immunohistochemical images of an identified VTA dopamine neuron locked in this plateau bursting mode are shown in Figures 4A and 4B, respectively. Figure 4A bottom trace illustrates that, in contrast to the rebound burst mode, the action potential threshold for plateau bursting showed a depolarizing shift of up to 10 mV before burst termination (Fig. 4A, bottom trace). As expected from this observation, the distribution of V_thr_ was clearly bimodal (Fig. 4D; V_thr_ BC: 0.66 ± 0.16; Mode 1 mean = −29.44 ± 4.83, SD = 1.16 ± 0.39; Mode 2 mean = −23.0 ± 8.12, SD = 2.42 ± 1.21; n = 6, N = 6), a unique feature of this type of bursting that is not seen in pacemaking or rebound bursting. Due to the occurrences of plateau potentials, the distribution of V_min_ for plateau bursts was complex, and could only be approximated with bimodal distribution (Fig. 4C; V_min_ BC: 0.53 ± 0.2; Mode 1 mean = −38.62 ± 5.0, SD = 2.81 ± 0.75; Mode 2 mean = −26.85 ± 6.49, SD = 1.18 ± 0.36, n = 6, N = 6). When plotting the ISI durations against V_thr_ (Fig. 4E), those V_thr_ residing in the more depolarized distribution (Mode 2 in Fig. 4D) were almost exclusively associated with very short intraburst ISI durations in the range of 6-26 ms (Fig. 4E, light green circles). Event-triggered averaging of action potentials associated with the V_thr_ Mode 2 distribution (Fig. 4F) revealed a distinct low-variability subthreshold membrane potential signature (Fig. 4F, bottom trace). What emerged was a sharply-rising membrane potential plateau of approximately 7 mV in amplitude and 30 ms in duration with a limited number of 2 to 3 high-frequency gamma-range spikes riding on top (plateau amplitude = 6.08 (5.22 to 7.87) mV; plateau duration = 35.78 (16.37 to 51.31) ms; number of spikes = 2 (2 to 2.14); mean maximum spike frequency = 53.26 (33.34 to 94.43) Hz). As evident from the average trace (Fig. 4F, bold trace), plateau burst intraburst frequency was less variable compared to intraburst frequencies in rebound bursting. Figures 4G and 4H show the firing frequency distribution of the example cell and the entire population of dopamine neurons identified as discharging in plateau burst mode, respectively. In essence, our unbiased exploration of membrane voltages from dopamine neurons *in vivo* uncovered two unique subthreshold burst signatures, each associated with a distinct and nonoverlapping frequency range.

**Figure 4.**
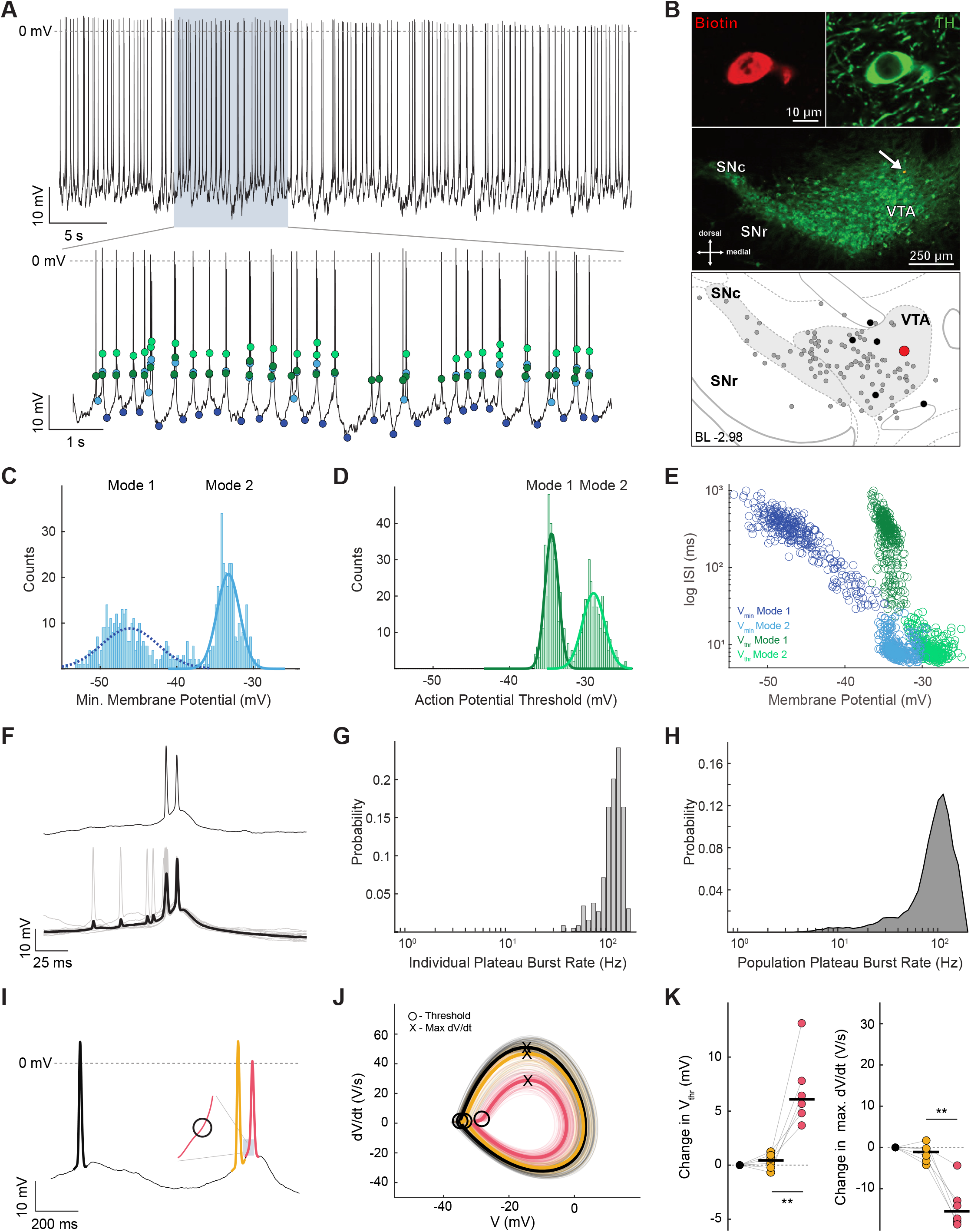
Plateau burst is characterized by a plateau depolarization and a depolarizing action potential threshold. A. Representative recording of a dopamine neuron in the VTA. The upper trace displays action potentials with overshoot during a stable recording for > 60 s. The higher temporal resolution in the lower trace allows visual identification of subthreshold events. Dark blue dots represent more hyperpolarized voltage minima whereas light blue dots represent minima occurring during plateaus. Dark green dots represent the subthreshold voltage maxima. Light green dots represent the more depolarized thresholds. B. Immunocytochemical identification (upper panel) demonstrates the recorded neuron (red) is TH-positive (green). Lower magnification image (middle panel) locates the recorded neuron (arrow) within the VTA. Depiction (bottom panel) shows the location of the recorded neuron (large red dot) in relation to all the recorded neurons in this study (gray and black dots). Black dots represent neurons that fired in a plateau burst pattern. C. Distribution of all subthreshold voltage minima (V_min_) from the cell in A-B, demonstrating a bimodal distribution (dark blue line = Mode 1 Gaussian fit; light blue line = Mode 2 Gaussian fit). D. Distribution of all subthreshold voltage maxima (V_thr_) from the cell in A-B, demonstrating a bimodal distribution (dark green line = Mode 1 Gaussian fit; light green line = Mode 2 Gaussian fit). E. Scatter plot of ISI versus V_min_ and V_thr_. The light green cluster is associated with more depolarized voltage membrane potentials and shorter duration ISIs. F. High temporal resolution of a plateau burst (upper panel), and event-triggered plateau burst average (black trace in lower panel) and overlaid (gray) traces. G. Probability histogram of the firing rate associated with plateau bursts for the example cell in A-B. H. Probability histogram of firing rate associated with plateau bursts for all cells. I. Example trace of a cell firing a plateau burst with three action potentials in order from black prior to the burst, then orange beginning the burst, and finally red ending the burst. Inset illustrates where the depolarized threshold occurs for the action potential ending the burst (red). J. Phase plots for each action potential are displayed in the same color scheme as in I. K. Summary data of the changes in threshold voltage (left) with significant depolarization in threshold of the action potential ending the burst (red), change in maximal depolarization rate (Max dV/dt; middle) with significant decrease in Max dV/dt of the last action potential (red) for all cells firing in a plateau burst pattern.

With regard to action potential trajectories, in contrast to the rebound burst, we found no change in the action potential threshold potential and waveform properties at the onset of the plateau burst (Fig. 4I-4K). This implies that no systematic and predictive threshold dynamics precede the occurrence of plateau bursts in dopamine neurons. Or stated from the neuron’s perspective, it was “surprised” by the burst. However, after the onset of plateau bursts, the threshold and action potential dynamics dramatically changed, as subsequent action potentials in this burst (in most cases the 2nd action potential) displayed a large depolarizing shift in the threshold potential associated with reduced depolarization speed, both indicators of reduced excitability (Fig. 4J and 4K; Δ V_thr_: spike 1 vs 2 = 0.39 (−0.41 to 0.92) mV, p = 0.36; spike 2 vs 3 = 6.13 (4.81 to 7.81), p = 0.002; Δ max dV/dt: spike 1 vs 2 = −1.15 (−3.25 to 0.12) V/s, p = 0.05; spike 2 vs 3: - 15.68 (−17.95 to −12.9) V/s, p = 0.0043; n = 6, N = 6; Supplemental Table 2). Thus, the threshold dynamics of the plateau burst not only occur in the opposite direction compared to those in rebound mode, but also serve to time the end, and not the beginning, of the burst. Indeed, in over 90 percent of all plateau bursts, the second action potential with the large depolarizing shift in the threshold was also the last action potential of the burst (plateau bursts with 2 spikes, n = 404; 3 spikes, n = 33; 4 spikes, n = 3). Additionally, a small amplitude spikelet was observed after the last action potential of the plateau burst (data not shown). Although dopamine neurons locked in the plateau burst mode were rare and only found in the VTA (Fig. 4B), a much larger percentage of dopamine neurons generated individual events in the gamma range (Fig. 1H). In fact, it is possible for dopamine neurons to fire in both rebound *and* plateau burst modes, as we have observed in a few neurons (data not shown).

In summary, the first systematic *in vivo* patch-clamp study of identified midbrain dopamine neurons provides direct evidence of two qualitatively distinct burst firing mechanisms characterized by contrasting subthreshold membrane potential signatures and opposing action potential threshold dynamics. These findings present the first framework of how predictive-coding might be biophysically implemented in midbrain dopamine neurons.

## Discussion

For the first time in nearly four decades, our study provides new intracellular *in vivo* recordings of identified midbrain dopamine neurons. By establishing deep *in vivo* wholecell patch-clamp recording with no noticeable perturbations on firing activity, our large dataset allowed for the first systematic analysis of the subthreshold membrane potential range of dopamine neurons in the intact brain. We demonstrate that *in vivo* pacemaker-like activity is associated with narrow unimodal distributions of the interspike subthreshold voltage minima and maxima, the latter representing the action potential thresholds. In contrast, transient accelerations in firing rate (i.e. bursting) were associated with two distinct subthreshold signatures and threshold dynamics: a prolonged membrane hyperpolarization leading to rebound bursts, which were associated with a hyperpolarizing shift in action potential threshold at the start of the burst; and a fast depolarization leading to plateau bursts, which were associated with a depolarizing shift in the action potential threshold at the end of the burst. While the plateau burst was strongly coupled to short gamma range discharge, the rebound burst enabled beta range bursts, but also allowed lower frequency discharges. This widening of the dynamic range of dopamine neuron firing by rebound bursting might be an attractive candidate mechanism to enable predictive coding for DA neurons *in vivo*. The hyperpolarizing shift of the action potential threshold induced in rebound burst mode also enhances the potential excitability of the neuron. At the same time, these dopamine neurons remain flexible to allow integration of incoming synaptic inputs. Moreover, the action potential threshold of the rebound burst spike is distinct from that of the background pacemaking, which would enable the cell to compute a new prediction for spike timing. Future *in vivo* whole-cell studies in awake behaving animals will be necessary to explicitly test whether this candidate subthreshold mechanism is indeed at work during ongoing reward prediction error computations. In contrast, the observed threshold dynamics of the short and high frequency plateau bursts are not likely to be utilized for predictive algorithms. Rather, they are reactive and might thus serve in controlling burst termination. These features render the plateau burst mechanism a plausible candidate for precise temporal signaling of, for example, unexpected and salient sensory events. While the subthreshold signature of the rebound burst suggests a timed-series of initial inhibition followed by excitation during the predictive time window, short plateau bursts might be generated by rapid fluctuations in the excitation/inhibition (E/I) balance. Indeed, our previous dynamic clamp studies *in vitro* revealed similar threshold and action potential waveform behavior when the E/I balance rapidly shifts toward more excitation or disinhibition (Lobb et al., 2011). In essence, our study identified *in vivo* operating principles of discharging dopamine neurons and provides explicit candidate mechanisms, which can now be tested in awake animals.

While the *in vivo* intracellular recordings with sharp microelectrodes of identified dopamine neurons from the 1980s did not provide systematic analyses of the subthreshold membrane potential range in relation to firing patterns, those results are mostly in accordance with our findings. There is, however, one important exception. Input resistances in our *in vivo* patch-clamp recordings of dopamine neurons are about 10-fold higher than those reported previously, indicating that the *in vivo* “high conductance” state was overestimated and might be in part caused by impaling dopamine neurons with sharp microelectrodes (Grace and Bunney, 1980, 1983a). In addition, it was now possible to determine that the *in vivo* whole-cell configuration did not alter firing patterns observed in intact cells – either with on-cell recordings in this study or by comparison to previous *in vivo* extracellular recordings of identified dopamine neurons (Farassat et al., 2019; Schiemann et al., 2012). Importantly, all neurons included were filled, recovered, and TH-immunopositive. Given these quality control measures and the large dataset, we can now begin to also address the *in vivo* subthreshold diversity of dopamine neurons. The previously reported differences in *in vitro* intrinsic subthreshold properties (Beier et al., 2015; Evans et al., 2017; Lammel et al., 2008; Neuhoff et al., 2002; Tarfa et al., 2017), such as sag amplitudes and rebound latencies, were also evident in our *in vivo* dataset, although dopamine neurons in the paranigral aspect of the VTA were underrepresented. These differences might therefore contribute to subpopulation-specific computations, as recently identified by *in vivo* GCaMP imaging during a complex behavioral task (Engelhard et al., 2019). Moreover, behaviorally-relevant somatodendritic computations of dopamine neurons relating to, for example, the current debate of coding motivation and learning (Mohebi et al., 2019) do not necessarily have to cross the action potential threshold (Alle and Geiger, 2006; Christie et al., 2011).

Despite these strengths, our study has a limitation in that we recorded under isoflurane anesthesia and were therefore unable to study subthreshold membrane potential signatures of dopamine neuron firing pattern in response to sensory cues and reward-related signals. However, the stability of brain states in controlled anesthesia is also an advantage as it enabled the unbiased identification of subthreshold membrane potential signatures associated with the two distinct burst discharge patterns. Another limitation of our study is that we did not go beyond the identification of subthreshold patterns to explicitly target underlying synaptic and intrinsic biophysical mechanisms. For instance, the deep hyperpolarization that precedes rebound bursting could be activated by different types of synaptic input involving distinct receptors ranging from GABA_A_, GABA_B_, dopamine receptor D2 (D2R) or metabotropic glutamate receptors coupled to small conductance calcium-activated potassium (SK) channels (Brazhnik et al., 2008; Fiorillo and Williams, 1998; Lacey et al., 1988; Lobb et al., 2011; Lobb et al., 2010; Paladini et al., 2001; Paladini and Tepper, 1999). In addition, the rebound kinetics are likely to be controlled by intrinsic conductances in the subthreshold range. Here, rebound depolarization accelerating (e.g. HCN channels; Neuhoff et al., 2002) and delaying (e.g. Kv4 channels; Subramaniam et al., 2014; Tarfa et al., 2017) conductances are relevant. Some DA neurons even respond to the termination of a hyperpolarizing current injection with a robust recruitment of T-type calcium channels (Evans et al., 2017; Wolfart and Roeper, 2002). The plateau burst, on the other hand, is likely driven by a rapid onset depolarization, which is likely to be caused by a rapid change in the E/I balance (Lobb et al., 2010). Fundamentally, the rebound and plateau bursts represent the responses within dopamine neurons that are comprised of a sequence of synaptic inputs that are active at any moment in the intact brain in addition to the intrinsic cellular processes that are particular to the individual neuron. With the use of our deep *in vivo* patch-clamp method, we unveiled the opposing biophysical mechanisms underlying the rebound and plateau burst signatures, providing the first framework for how coding might be biophysically implemented in midbrain dopamine neurons. This method will facilitate future mechanistic studies of dopamine neurons in awake animals, as it enables not only the control of intracellular milieu (e.g. calcium buffering) but also the direct intracellular application of pharmacological agents, for studying behaviorally-relevant dopamine neuron firing.

## Materials & Methods

### Animals

C57BL/6N (Charles River Laboratories and Janvier Labs) and C57BL/6J (Jackson Laboratories) mice (67 males and 7 females, aged 8-16 weeks) were used for the experiments. Mice were maintained on a 12-hour light/dark cycle with food and water available *ad libitum*. All experimental procedures were approved by the German Regional Council of Darmstadt (TVA 54-19c20/15-F40/28) or the University of Texas at San Antonio Institutional Animal Care and Use Committees.

### Stereotactic surgeries

For head-plate implantations, animals were anaesthetized with isoflurane (1.0-2.5% in 100% O2, 0.35 L/min) and placed in a stereotaxic frame (David Kopf Instruments). The body temperature was maintained at 37-38 °C with a heating blanket, and the breathing rate at 1-2 Hz. Small indentations were made above the VTA (AP: −3.08 mm, ML: ±0.25-0.75 mm) or SNc (AP: −3.08 mm, ML: 0.9-1.4 mm) as a reference for later craniotomies. Customized head-plates were anchored to the skull with superglue and dental cement (Paladur, Kulzer), as well as with screws mounted on the skull.

### In vivo electrophysiology

Electrophysiology was performed using glass electrodes (5-10 MΩ) filled with internal solution containing (in mM): 135 K-gluconate, 10 HEPES, 5 KCl, 5 MgCl2, 0.1 EGTA, 0.075 CaCl2 (variant 1) or no added CaCl2 (variant 2), 5 ATP (Na), and 1 GTP (Li), neurobiotin (0.1%; Vector Laboratories) or biocytin (3%), at pH 7.35. The estimated free calcium concentrations based on a calcium contamination of 15 μm (for details see Woehler et al., 2014) were about 40 nM for variant 1 and about 80 nM for variant 2. Recording electrodes were made with borosilicate glass capillaries (G120F-4, Warner Instruments) pulled with a pipette puller (DMZ Universal Electrode Puller, Zeitz-Instruments).

Animals were maintained under isoflurane (1.0-2.5% in 100% O2, 0.35 L/min) and head-fixed to a customized recording platform (Luigs & Neumann). The body temperature (37-38 °C) and breathing rate (~1 Hz) were continuously monitored. Craniotomies and duratomies were performed at the reference locations. The electrode was lowered down to approximately 200 μm above the region of interest with a positive pressure of 600-1000 mbar. The pressure was gradually lowered and kept in a range between 40-70 mbar during probing for cells in the VTA (DV: 4.0-5.2 mm) and SNc (DV: 3.8-5.2 mm). A tone generator (PSA-12, HEKA; Custom script, Axograph) was used to monitor the pipette resistance during probing and sealing. A fluctuating 20-50% increase in the pipette resistance indicated a cell was nearby. Once reaching proximity, pressure was released to achieve a GΩ seal. On-cell recordings were obtained for some neurons prior to breaking in. After breaking into the whole-cell mode with a small suction, cell capacitance and series resistance were estimated. Whole-cell recordings of spontaneous activity and responses to hyperpolarizing current injections were then obtained in the current-clamp mode. Recording signals were obtained with a patch-clamp amplifier (EPC10 USB, HEKA; Multiclamp 700B, Molecular Devices) and the data were acquired with PatchMaster software (HEKA) or Axograph software (Axograph) at a sampling rate of 20 kHz.

### Histology and microscopy

After recording, animals were injected with a lethal dose of pentobarbital (0.3-0.4 mL) and transcardially-perfused with PBS followed by 4% paraformaldehyde (wt/vol) and 15% picric acid (vol/vol) in phosphate-buffered saline. Collected brains were post-fixed overnight. Immunocytochemistry and confocal microscopy were performed as previously described (Schiemann et al., 2012). For the tyrosine hydroxylase (TH) staining, polyclonal rabbit antibody (catalog no. 657012, Merck) was used as the primary antibody, and AlexaFluor-488 goat anti-rabbit IgG (catalog no. A11008, ThermoFisher) as the secondary antibody. Neurobiotin and biocytin were visualized with AlexaFluor-568 streptavidin conjugate (catalog no. S11226, ThermoFisher).

### Spike detection

A spike peak was detected if the following three conditions were met within a 3 ms window after crossing a dV/dt threshold (default value = 10 mV/ms): First, the minimal value of dV/dt must be less than −5 mV/ms. Second, the maximal value of voltage must be within 30 mV of the highest voltage in the entire trace. Last, the difference between the maximal voltage and the voltage at the dV/dt-threshold crossing (spike height) must be greater than 5 mV. In some cases, these parameters were modified to improve spike detection performance.

### Quantification and statistical analysis

Data processing and quantifications were performed using MATLAB (MathWorks). Wilcoxon Rank sums tests were used to determine statistical differences using MATLAB (MathWorks). Graphs and maps were created using MATLAB or Prism 6. Bimodality coefficient was used for dual process detection (Freeman and Dale, 2013). Measurements of effect size were also calculated (Hentschke and Stuttgen, 2011). Parametric data were presented as mean ± standard deviation. Non-parametric data were presented as median with interquartile range.

### Data availability

Access to all original patch-clamp data files, immunohistochemical and anatomical data of identified dopamine neurons, as well as MATLAB scripts for analysis, will be provided upon full publication.

## Acknowledgements

We are grateful to Nao Uchida, who helped define the relevant *in vivo* patch-clamp recording parameters of dopamine neurons. We thank Kauê M. Costa for his help with confocal imaging and initial mapping of dopamine neurons. We are grateful for excellent technical support by Jasmine Sonntag and Beatrice Fischer (Neurophysiology, Goethe University Frankfurt). We thank Gerard Beaudoin for custom scripting of Axograph tone generation. We thank Stephan Lammel (UC Berkeley) for his advice on the manuscript and Julia Kuhl (http://www.somedonkey.com) on the design of the figures. This work was funded by grants to JR (DFG CRC1080, CRC1193 & NIH DA041705) and CP (NIH MH113341, MH107229).

## Author Information

JR & CP designed the study and established the technique for *in vivo* patch-clamp recordings of identified dopamine neurons during CP’s sabbatical in Frankfurt. KO, JP & CP performed the experiments. Analysis was carried out jointly by KO & SS supervised by JR, and AK & JP supervised by CP.

## Ethics Declaration

Authors declare no conflict of interests

---

**Supplemental Figure 1.**
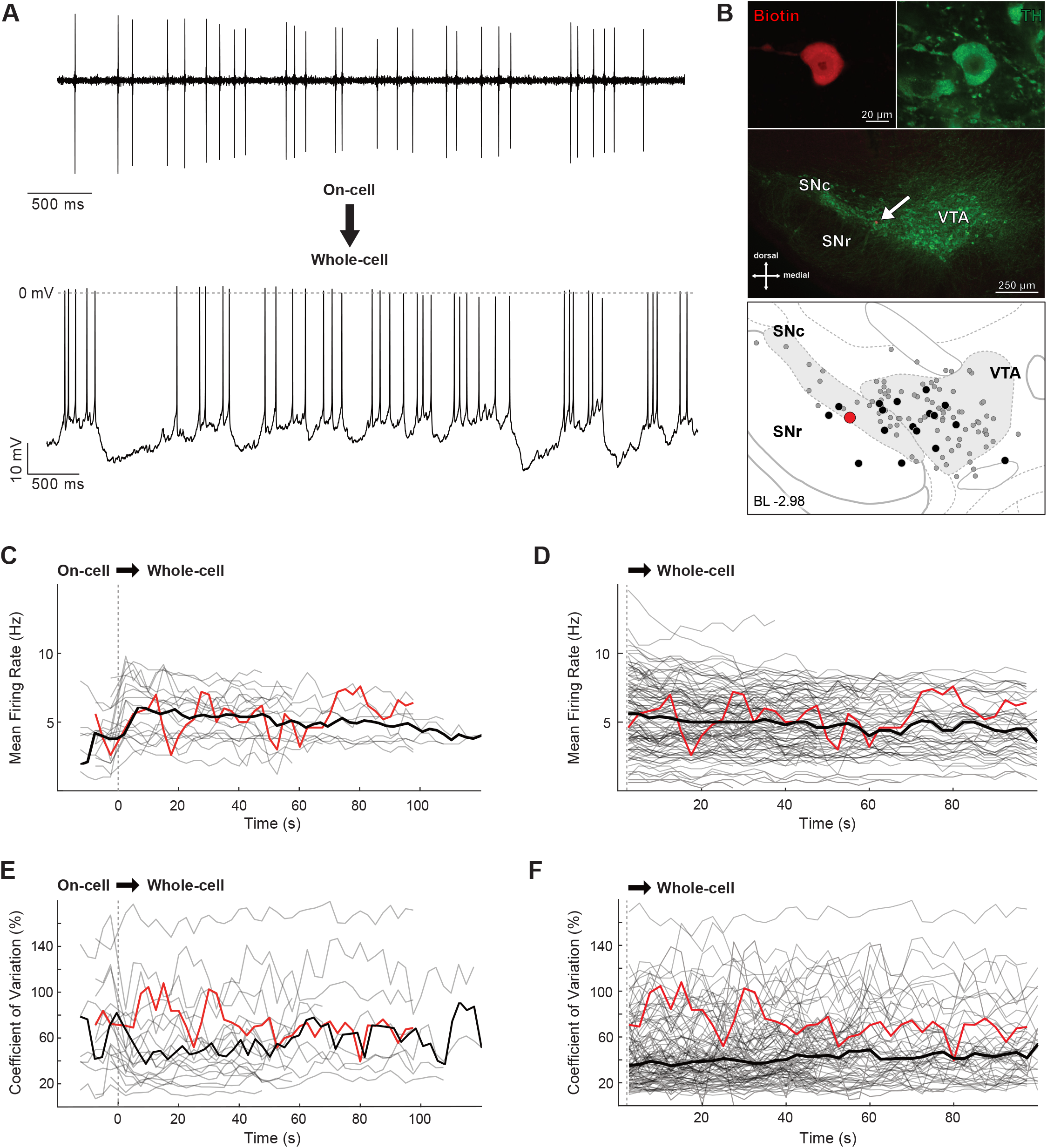
*In vivo* whole-cell recording of dopamine neurons does not disturb firing rate or pattern of activity. A. Representative recording of spontaneous *in vivo* electrical activity from a neuron recorded in on-cell mode (upper trace) and then broken into the whole-cell configuration (lower trace) with similar firing rate and pattern. B. Immunocytochemical identification (top panel) demonstrates the recorded, biocytin-filled neuron (B; red) is positive for tyrosine hydroxylase (TH; green). Lower magnification image (middle panel) locates the recorded neuron (arrow) within the medial region of the SNc. The depiction (lower panel) shows the location of the recorded neuron (large red dot) in relation to all the recorded neurons in this study (black and gray dots). Black dots represent all cells for which on-cell activity was recorded for comparison with whole-cell activity. C. Time series of mean firing rate during the transition from on-cell to whole-cell modes (vertical dashed line at time = 0; on-cell mean rate = 4.27 (3.18 to 5.75) Hz; wholecell mean-rate = 5.76 (3.92 to 6.56) Hz; n = 19, N = 16; p = 0.11). D. Time series of mean firing rate for the first 100 seconds from all neurons recorded in this study (slope change in FR = −0.005; n = 110, N = 78). E. Time series of the coefficient of variation (CV) of firing during the transition from on-cell to whole-cell mode (vertical dashed line at time = 0; on-cell CV = 0.69 (0.42 to 0.82); whole-cell CV = 0.51 (0.34 to 0.77); n = 19, N =16; p = 0.38). F. Time series of firing CV for the first 100 seconds from all neurons recorded in this study (slope change in CV = 0.003; n = 110, N = 78). C-F. Red lines represent the example neuron illustrated in A and B. Black lines represent the average of all lines for each panel.

**Supplemental Figure 2.**
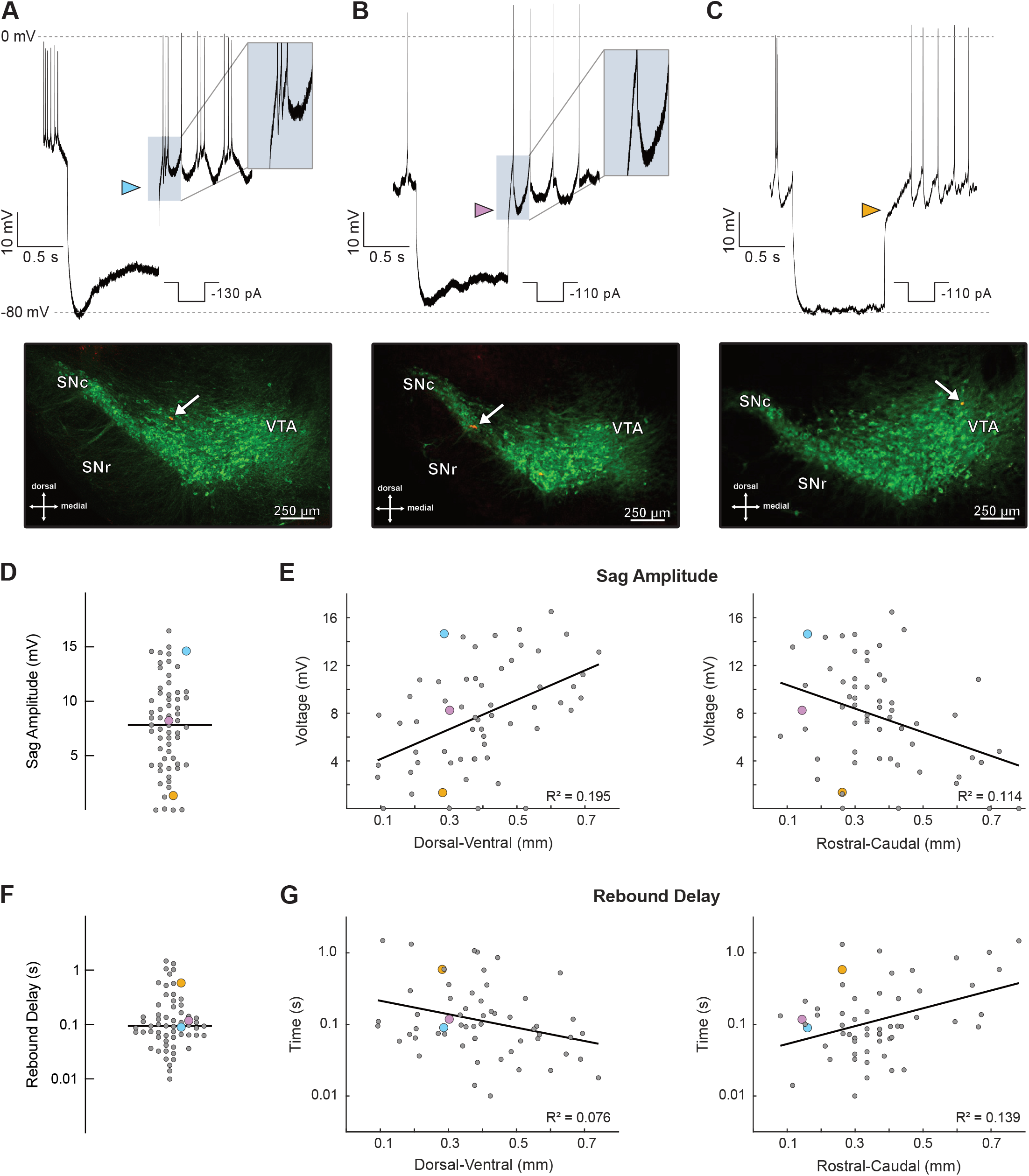
Sag and rebound delay are correlated with cell location. A. Example recording from an identified dopamine neuron with a large sag in response to hyperpolarizing current injection, and a short delay to the first spike upon removal of the hyperpolarization (blue arrowhead). A-C. The immunocytochemical image shows the location (arrow) of the recorded neuron. B. Example recording from an identified dopamine neuron with a moderate sag in response to hyperpolarizing current injection, and a moderate delay to the first spike upon removal of the hyperpolarization (pink arrowhead). C. Example recording from an identified dopamine neuron with almost no sag in response to hyperpolarizing current injection, and a long delay to the first spike upon removal of the hyperpolarization (orange arrowhead). D,F. Plots of the sag amplitude (D) and rebound delay (F) for all neurons where the relevant protocols were carried out (sag amplitude at −80 mV = 8.16 ± 4.41 mV; n = 60, N = 50; rebound delay = 84.55 (52.84 to 146.44) ms; n = 60, N = 50). E,G. Relationship of cell location with sag amplitude and rebound delay. Both measures had a significant relationship with location in the dorsal-ventral and rostral-caudal planes. D-G. Large dots are colored in correspondence with the arrowhead colors in A-C.

**Supplemental Data Table 1.** Table showing that there were no significant differences in firing rate and CV of ISI between various groupings of cells:

| Comparison type | FR (Hz) |  |  | CV |  |  |
| --- | --- | --- | --- | --- | --- | --- |
|  | N | J | p-value | N | J | p-value |
| **C57BL/6N (n=92, N=65) vs C57BL/6J (n=20, N=15)** |  |  |  |  |  |  |
|  | 5.09<br>(3.46-6.8) | 5.39<br>(3.61-6.62) | 0.94 | 0.41 (0.26-0.62) | 0.49<br>(0.39-0.68) | 0.15 |
| **Male (n=93, N=67) vs Female (n=12, N=7)** | m | f | p-value | m | f | p-value |
|  | 5.23<br>(3.65-6.72) | 6.07<br>(4.38-7.2) | 0.31 | 0.43 (0.3-0.67) | 0.45<br>(0.41-0.58) | 0.45 |
| **[Ca<sup>++</sup>] 40nM (n=8, N=6) vs 80nM (n=104, N=74)** | 40 | 80 | p-value | 40 | 80 | p-value |
|  | 3.86 (1.0-6.12) | 5.26<br>(3.59-6.8) | 0.15 | 0.45 (0.39-0.66) | 0.42 (0.3-0.65) | 0.71 |

- Cell from animals belonging to sub-strains C57BL/6N vs C57BL/6J
- Cells from male vs female animals
- Recordings made using internal solution containing 40 nM vs 80 nM free calcium All values are reported as median (25th quantile-75th quantile), and all p-values are reported for the Wilcoxon rank sum test. Upper-case letter ‘N’ indicates the number of animals and the lower-case letter ‘n’ indicates the number of cells in that grouping.

**Supplemental Data Table 2.** Table showing trajectories of repolarization speed (min dV/dt) across three types of spiking activity patterns. All values are reported as median (25th quantile-75th quantile), and all p-values are reported for the Wilcoxon rank sum test.

| Spiking pattern | min dV/dt spike 1 vs spike 2 (V/s) |  | min dV/dt spike 2 vs spike 3 (V/s) |  |
| --- | --- | --- | --- | --- |
|  | Magnitude | p-value | Magnitude | p-value |
| Pacemaker | 0.03 (-0.09 to 0.21) | 0.78 | -0.0 (-0.16 to 0.08) | 0.45 |
| Rebound burst | -3.6 (-6.85 to -2.21) | $1.15 \times 10^{-7}$ | 2.03 (1.26 to 2.86) | $7.05 \times 10^{-7}$ |
| Plateau burst | 0.94 (-0.67 to 1.84) | 0.68 | 7.35 (6.66 to 9.68) | 0.008 |

## Notes

### Competing Interest Statement

The authors have declared no competing interest.

